# Detecting Temporal Changes in Skin Surrogate Model Using Hyperspectral Imaging

**DOI:** 10.1101/2025.03.20.644453

**Authors:** Neetu Sigger, Tuan T. Nguyen, Gianluca Tozzi

## Abstract

Advancements in biomedical imaging have increasingly explored innovative models for studying human skin conditions. Recent research highlights a biochemical connection between banana peels and human skin, particularly through the enzyme tyrosinase, which plays a key role in pigmentation and oxidative stress. Studies have shown that, much like human skin, banana peels undergo oxidative browning due to tyrosinase activity. Leveraging this similarity, we have developed imaging techniques to track biochemical changes in bananas as a potential surrogate model for the detection of skin changes such as sun spots or melanoma. This paper introduces the application of hyperspectral imaging (HSI) to analyse the spectral properties of banana peels as a method for studying pigmentation disorders and oxidative damage. We present a HSI system designed to capture spectral reflectance characteristics across the 440 nm to 900 nm range, with a hyperspectral cube resolution of 956×952×96. Our approach enables a more targeted spectral assessment by dynamically selecting and investigating specific regions in banana samples over multiple days. Preliminary spectral analysis demonstrates the system’s capability to detect reflectance variations linked to biochemical changes such as ripeness, bruising, and oxidative stress—factors that parallel pigmentation alterations in human skin.

However, direct hyperspectral imaging can be resource intensive, making it less accessible for applications in dermatology and mobile health diagnostics. To address this, we conducted preliminary experiments to reconstruct 27-band hyperspectral representations in the VIS spectrum from RGB images. The observed spectral variations and successful reconstruction results confirm the potential of HSI for capturing biochemical changes and provide a foundation for future biomedical imaging research, particularly in melanoma detection and dermatology-related applications.

## Introduction

Medical imaging has revolutionised dermatological diagnostics, enabling early detection of conditions such as melanoma, a highly aggressive form of skin cancer [24, 25]. Traditional diagnostic techniques, including dermoscopy and histopathology, remain the gold standard; however, these methods are often subjective and time-intensive [1, 30, 33, 4]. To address these challenges, non-invasive imaging methods such as hyperspectral imaging (HSI) have been explored, providing detailed spectral information beyond the visible range and enabling precise biochemical tissue characterisation [24, 15, 38]. HSI is a powerful imaging modality that captures and analyses spectral data across multiple wavelengths, spanning the visible to near-infrared (NIR) spectrum [16, 21]. Unlike conventional imaging techniques, HSI identifies unique spectral signatures related to tissue composition, oxidative stress, and pigmentation anomalies, which makes it particularly relevant for skin cancer diagnosis [14, 11]. Recent studies have demonstrated the effectiveness of HSI in detecting melanoma by analysing subtle spectral variations linked to tyrosinase activity and abnormal pigmentation [13, 2]. Furthermore, HSI enables real-time, non-invasive monitoring, which enhances diagnostic accuracy and facilitates disease progression tracking.

Despite these advantages, certain challenges must be addressed for the widespread application of HSI in dermatology [10, 9]. The availability of HS skin databases is currently limited, with existing datasets focused on facial recognition or skin colour modelling rather than medical diagnostics [7, 34]. In addition, ethical considerations play a crucial role in human skin imaging, which requires strict adherence to protocols for informed consent, data privacy, and compliance with biomedical research guidelines [3, 23, 8]. These ethical restrictions limit the collection of large-scale hyperspectral datasets from human subjects and further slow the progress in dermatological applications. Another challenge in acquiring hyperspectral skin data is the extended duration required to monitor changes in human skin. Since many dermatological conditions develop gradually, the data collection process becomes increasingly time-intensive.

To address these challenges, recent studies have explored the use of synthetic skin phantoms and plant-based alternatives, such as banana peels, as cost-effective and ethically viable surrogate models to simulate the properties of human skin [22]. Both human skin and banana peels contain tyrosinase, an enzyme essential for pigmentation and oxidative stress responses [32, 18]. In humans, tyrosinase plays a critical role in melanin production, whereas, in bananas, it regulates oxidative browning, a process with notable biochemical parallels to changes in skin pigmentation [18, 26]. Using this similarity, banana peels provide a cost-effective and ethically viable alternative for the acquisition of hyperspectral data. We used banana peels as a skin surrogate model for our initial experiments, allowing spectral analysis in an indoor setting while reducing the ethical and logistical barriers associated with direct human skin imaging.

However, direct HSI acquisition can be resource-intensive, limiting its accessibility for dynamic or real-time settings. This limitation restricts the broader application of HSI, especially in scenarios requiring rapid spectral analysis. To overcome this challenge, various methods have been explored, including snapshot compressive imaging systems and computational reconstruction algorithms, which convert 2D measurements into 3D hyperspectral cubes [5, 37]. Although promising, these approaches often depend on high-cost hardware, making them impractical for widespread implementation. To reduce dependence on costly hyperspectral imaging systems, spectral reconstruction algorithms have been developed to generate hyperspectral data from RGB images [6, 28]. These techniques seek to extract spectral details beyond the visible range, providing a more suitable alternative to direct HSI acquisition.

The aim of this study is to explore the potential of HSI for analysing biochemical and spectral variations using a skin surrogate model. Specifically, we (1) captured hyperspectral reflectance data from banana peels to simulate skin conditions such as melanoma and sunspots; (2) tracked temporal and spatial spectral variations over multiple days through dynamic region-of-interest (ROI) selection; and (3) extracted RGB images from hyperspectral data and employ AI models to reconstruct 27-band hyperspectral representations in the visible spectrum, enhancing spectral reconstruction for biomedical imaging applications.

## Method and Data collection

This section describes the methodology used for data collection and presents a detailed analysis of the captured dataset. We then discuss the evaluation metrics for RGB-to-HSI mapping, ensuring accurate representation across both spatial and spectral domains.

### Spectral and Spatial properties

Figure 1 illustrates the Living Optics HSI camera, which supports both scan-based and snapshot-imaging methods for acquiring HSIs. The banana dataset was captured using the snapshot imaging method, which includes a wavelength range of 440 nm to 900 nm, covering both the visible and nearinfrared spectrum. The system records data across 96 spectral channels, providing sufficient spectral resolution to detect subtle variations in surface reflectance that are due to changes in biochemical composition. This acquisition method captures the entire hyperspectral cube in a single exposure, eliminating motion artefacts.

**Figure 1:**
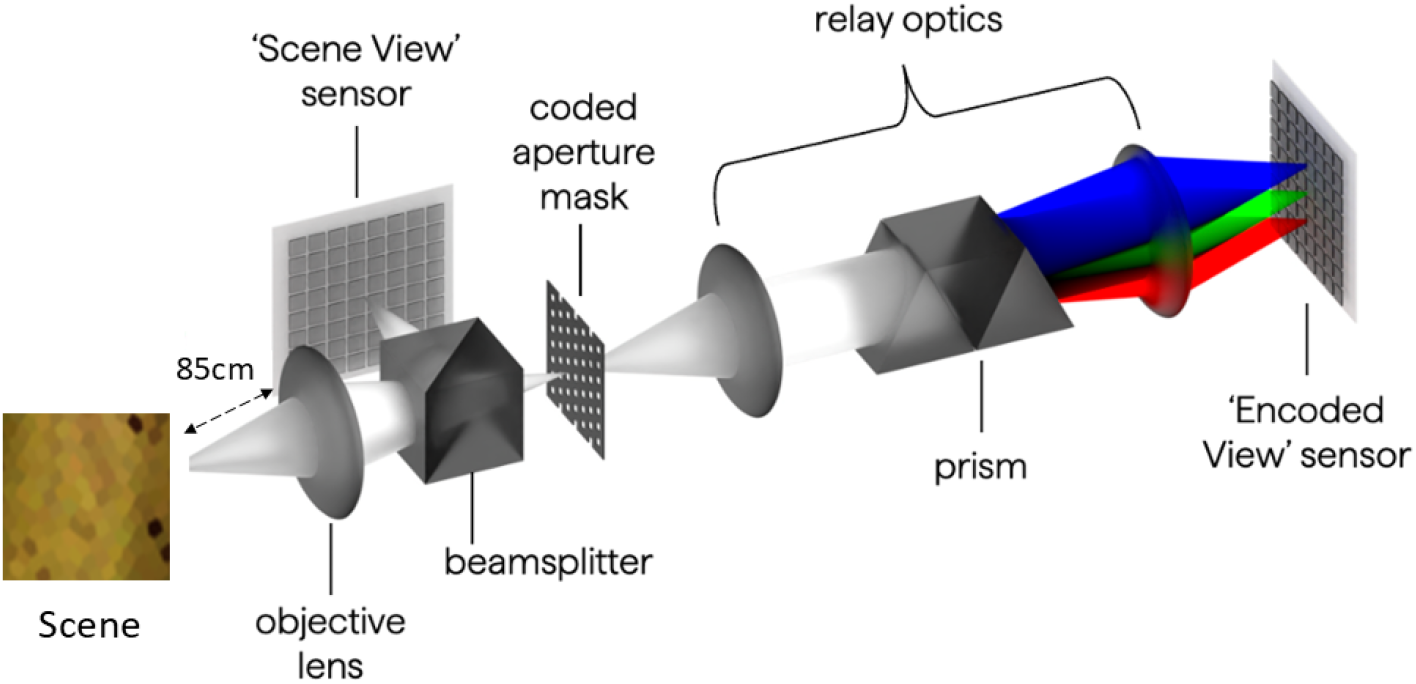
The Living Optics camera [31]. Light from the scene enters through the objective lens, where a portion is directed onto the ‘Scene View’ sensor. The remaining light passes through a coded aperture mask containing 4384 apertures, each corresponding to a discrete sampling point in the field of view. The transmitted light is then relayed through an optical system with a dispersive element, projecting 4383 dispersed beams onto the ‘Encoded View’ sensor. The camera software computationally decodes this image, producing 96 spectral channels for each sampling point.

A crucial consideration for the HSI system designed for banana skin analysis is the spatial resolution and field of view (FoV). The system records 4,384 evenly distributed sampling points with a spatial resolution of 956 × 952 pixels. These points are arranged in a 16 × 274 grid, with each sampling point covering a 3 × 3 pixel area, corresponding to 8.22 *µ*m^2^ on the sensor. The alignment between spatial resolution and FoV ensures accurate imaging throughout the banana surface while maintaining portability and ease of use.

### Data Collection

The distance between the HSI system and the target area was fixed at 85 cm for all samples as shown in Fig. 1. It was controlled using a computer running the Living Optics Software Development Kit (SDK) ^1^. The system operated at a frame rate of 200 mHz, with LED lighting used to illuminate the scene for consistent spectral acquisition.

HS images were collected over a seven-day period from banana samples to monitor spectral variations associated with pigmentation changes and structural transformations. To ensure precise spectral measurements, a calibration step calibration step was performed before data processing. For reflectance calculation, raw HS images *X* were corrected using a white reference *W* and a dark reference *D*. The dark reference image *D* was obtained by closing the camera lens during capture and deactivating the light source, ensuring that no external illumination influenced the measurements. Additionally, a white reference *W* was recorded using a Tyvek sheet [19]. To improve accuracy, five recordings were taken for both dark *D* and white *W* references, and the mean value was used for the reflectance calculation.

The reflectance (*R*) was then calculated using the following equation:

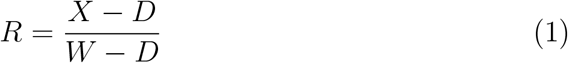

The acquired HS image contains background information that does not provide useful data for spectral analysis, potentially degrading the performance of the HSI system due to the simultaneous capture of background noise and unwanted artefacts [36]. To mitigate these effects, we selected ROIs in the acquired images, ensuring that only relevant spectral data are analysed while minimising interference from background and non-informative pixels. For each acquired HS image, six ROIs per image were selected to systematically track temporal and spatial spectral variations, as shown in Figure 2(a). Each selected ROI was cropped to 128 × 128 pixels to establish the dataset. The dataset includes HS images and spectral reflectance data corresponding to each ROI (Figure 2(b)), providing a structured approach to analysing spectral variations over time.

**Figure 2:**
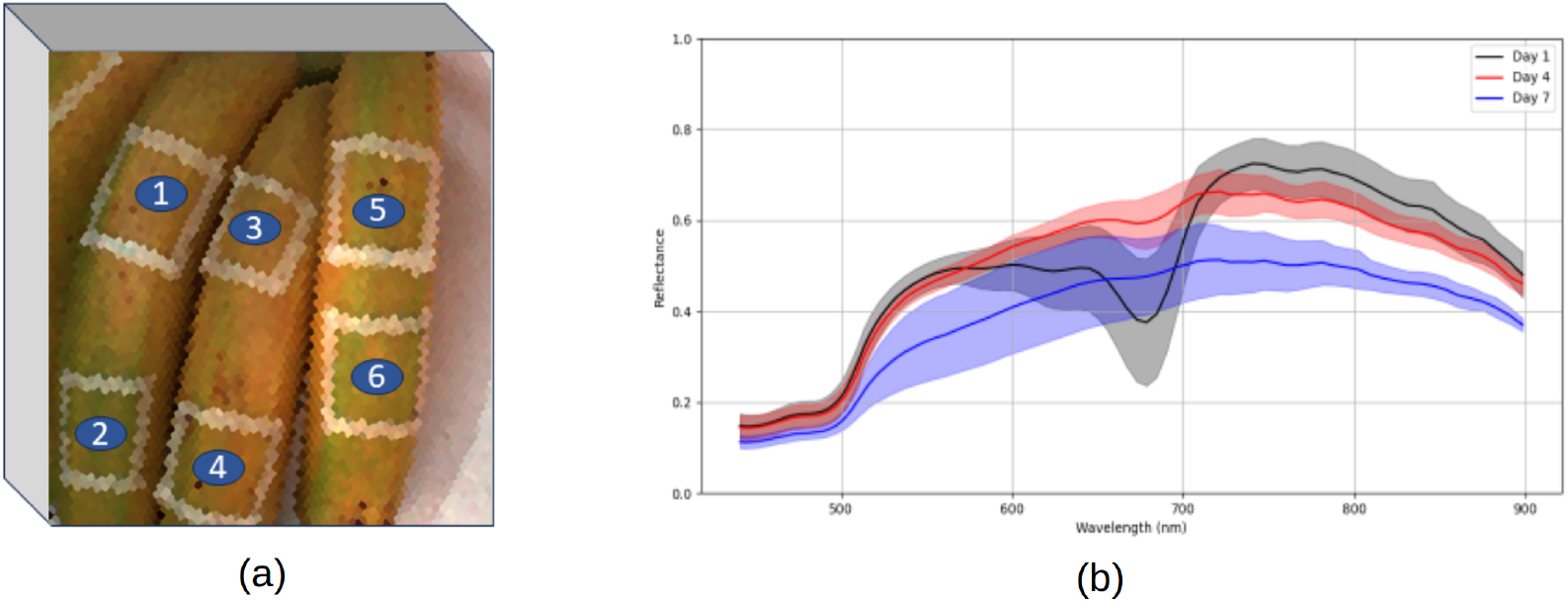
(a) Locations of the six selected regions of interest (ROIs) on HSI. (b) Average spectral reflectance of six ROIs (solid lines) with standard deviation (shaded areas) for Day 1, Day 4, and Day 7.

### Evaluation Metrics

To assess the quality of the reconstructed hyperspectral data, we utilise two evaluation metrics: Structural Similarity Index (SSIM) [39] for evaluating spatial accuracy and Spectral Angle Mapper (SAM) [29] for assessing spectral consistency.

Let *H* ∈ ℝ ^*w×h×C*^ be the original hyperspectral cube and 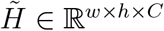 be the reconstructed version, where *C* = 27 for our dataset.

For spatial evaluation, SSIM is used to compare the structural similarity between ROIs of the original and reconstructed images at each spectral band. Given ground truth and reconstructed ROIs, 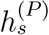 and 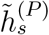, SSIM is formulated as:

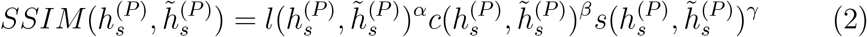

where *l, c*, and *s* represent luminance, contrast, and structural components, respectively, and *α, β, γ* are their corresponding weighting factors.

For spectral evaluation, SAM determines the spectral fidelity by computing the angular difference between the spectral signatures of the reconstructed and ground truth cubes. The SAM metric for a pixel (*i, j*) is given by:

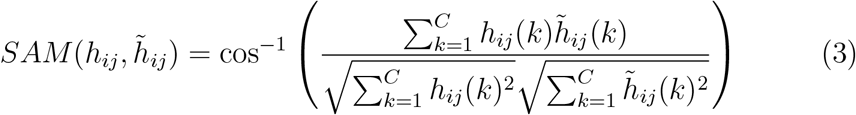

where *h*_*ij*_(*k*) and 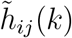 denote spectral intensity values in band *k* for the original and reconstructed hyperspectral cubes, respectively. SAM values range from 0 to *π*, where a lower SAM value indicates a closer spectral match.

## Results and Discussion

To evaluate the suitability of the HSI camera for its intended application, a preliminary analysis of the acquired samples was conducted. The objective was twofold: (1) to analyse the spectral signatures extracted from the collected HS images, ensuring their relevance for detecting the changes in skin model, and (2) to improve accessibility by developing an efficient RGB-to-HSI mapping, enabling broader applications in biomedical imaging and skin research.

### Analysis of Spectral Reflectance

The collected hyperspectral data was processed using Python and the Living Optics SDK. HS reflectance data obtained from six ROIs across multiple days reveal key biochemical transformations occurring during banana ripening and oxidation. As bananas ripen, their chemical composition changes, leading to modifications in reflectance spectra. One of the most significant indicators of ripening is chlorophyll degradation, which directly affects the spectral properties of the peel [17, 27]. The mean spectral reflectance of the six ROIs is illustrated in Figure 2(b). Initially, in the early ripening stage (day 1), the banana peel exhibits a high chlorophyll content, giving it a green appearance. This is reflected in a strong absorption trough near 680 nm in the visible (VIS) spectrum (Figure 2(b)), characteristic of chlorophyll absorption. At this stage, the reflectance in the NIR region (700–900 nm) remains high, indicating the presence of moisture content and structured pigmentation. On day 4, as the banana ripens, the chlorophyll content decreases, causing a shift in peel colour from green to yellow. This phase is marked by a progressive reduction in the chlorophyll absorption feature at 680 nm, while the overall reflectance spectrum in the 550–900 nm range becomes smoother. This spectral transition is related to the breakdown of chlorophyll molecules and the accumulation of carotenoids, which contribute to the characteristic yellow hue of ripe bananas. In the final stages of ripening (Day 7), the banana peel undergoes enzymatic browning, which further alters its spectral properties. The reflectance spectrum becomes progressively flattened, indicating severe oxidative stress and pigmentation loss (Figure 2). This trend aligns with previous studies [20], which demonstrate that as bananas approach the end of their ripeness cycle, their spectral reflectance curves lose distinct absorption features, resulting in a smooth spectral signature.

Furthermore, we focus on specific ROIs rather than analysing the entire banana peel to ensure a targeted and high-contrast spectral assessment of biochemical transformations. Distinct spectral trends were observed in different ROIs (Figure 3), revealing localised differences in ripening progression. ROI 1 (Fig. 3a) and ROI 3 (Fig. 3c) exhibited rapid declines in reflectance (700–900 nm), suggesting significant moisture loss and enzyme browning, likely corresponding to areas of early oxidation and browning spots. In contrast, ROI 2, ROI 4 and ROI 5 showed a moderate decline in reflectance, indicating progressive pigment breakdown and yellowing, as illustrated in Figures 3c, 3e, 3f, respectively. These regions appear to be in an intermediate ripening state, with some chlorophyll content still intact. Meanwhile, ROI 6 retained a higher NIR reflectance, implying delayed oxidation and higher moisture retention, likely due to reduced exposure to oxidative factors. Figure 4 also consolidates these spectral trends, demonstrating how different regions undergo distinct biochemical transformations over time.

**Figure 3:**
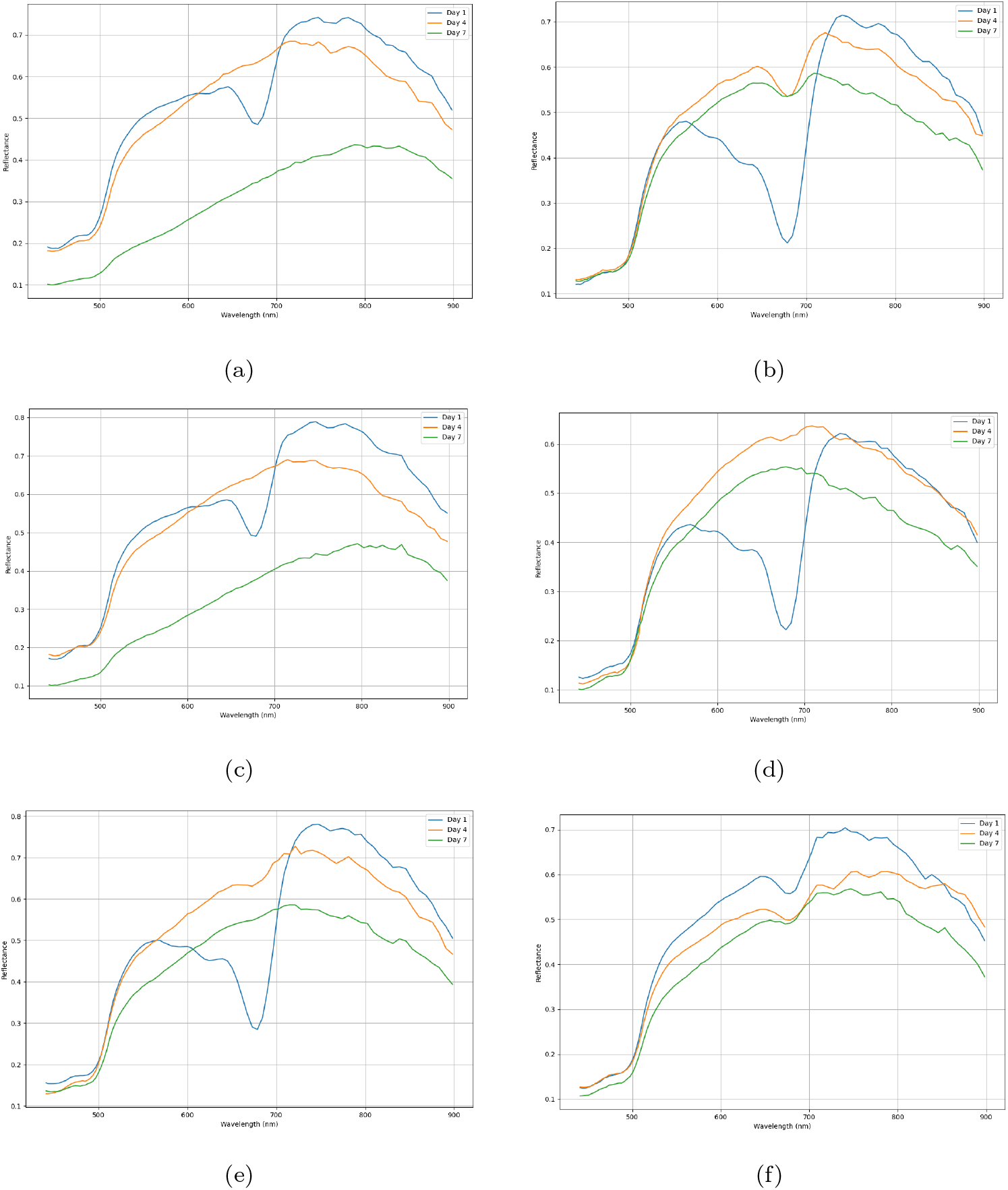
Spectral reflectance obtained from each ROI1(a)-ROI6 (f) for Day 1, 4, and 7.

**Figure 4:**
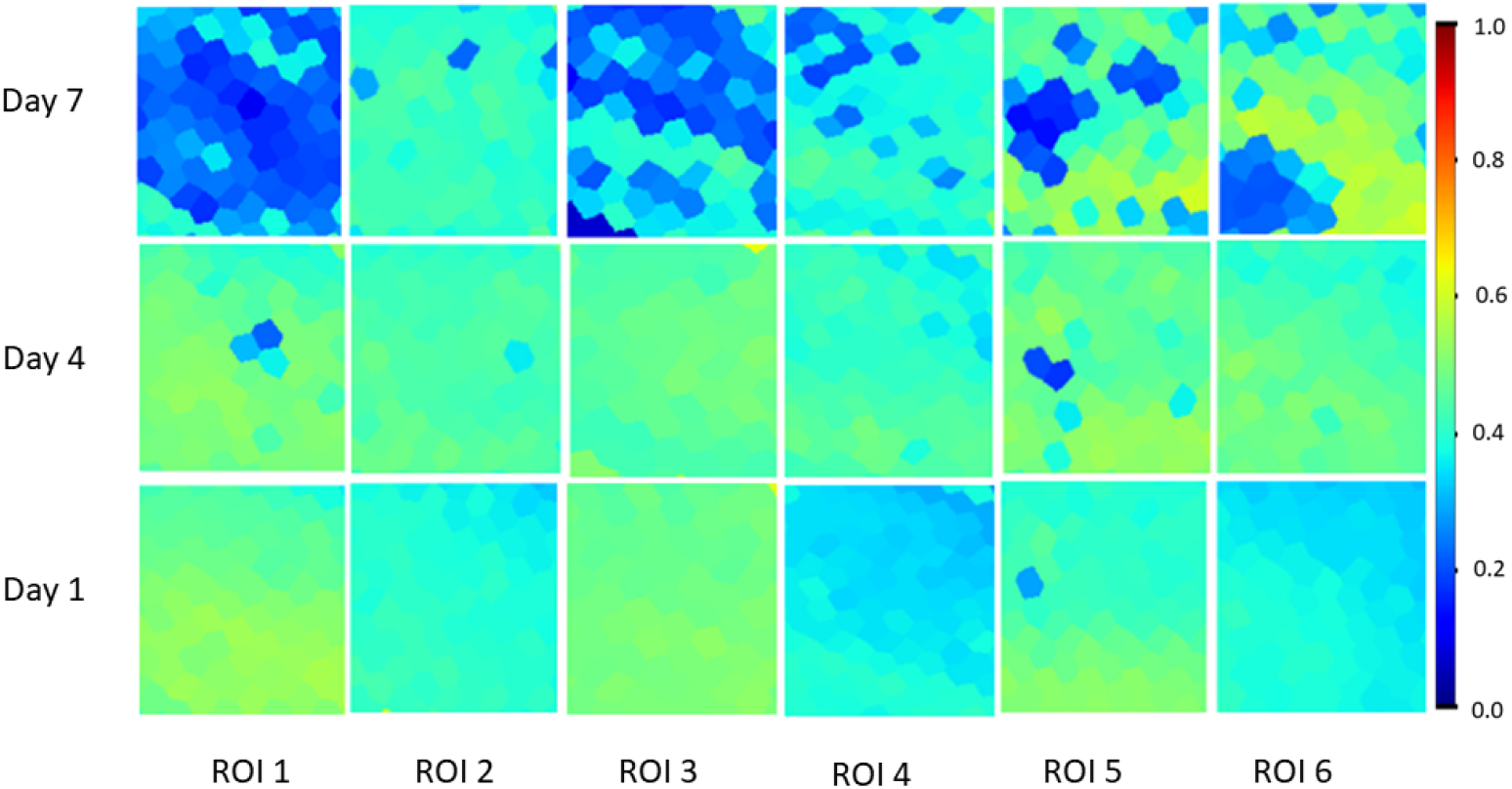
Spectral changes expressed as normalised reflectance in the ROIs (1-6) over time, shown for Day 1, 4, and 7.

We observed a decline in reflectance near 680 nm, which indicates chlorophyll breakdown, while the flattening of the spectral characteristics in the VIS and NIR ranges is associated with melanin accumulation and oxidative browning [35, 12]. Localised variations across different ROIs highlight the complexity of biochemical changes, and the findings demonstrate the potential of HSI for precise spectral characterisation of skin-like materials.

## Spectral Reconstruction from RGB

Various approaches have been proposed for HS reconstruction from RGB images. Readers seeking a comprehensive overview of these methods can refer to the survey by [5], which summarises key models developed over the past two decades. For our benchmark design, we specifically selected the Multi-stage Spectral-wise Transformer (MST++) [6] model due to its lower computational cost and faster processing time, making it a suitable choice for real-time biomedical imaging applications.

Figure 5 illustrates the results using the pre-trained MST++ model, which was applied to the RGB-to-HSI mapping. Since MST++ was trained on the NTIRE dataset (400-700 nm), we discarded the first four channels in the reconstruction results to align with our dataset (440-900 nm). Channels beyond 700 nm, unsupported by the pre-trained model, were excluded from evaluation. Table 1 presents the spectral and spatial evaluation results using SAM and SSIM for the six ROIs in our dataset. In our dataset, ROI 1, ROI 3, and ROI 5 exhibited relatively lower SAM values, suggesting greater spectral deviations due to oxidative stress and pigment breakdown. In contrast, ROI 2 had the highest SAM value (0.610), suggesting a slightly higher deviation from the expected spectral signature. In particular, ROI 3 achieved the highest SSIM (0.596), demonstrating strong spatial preservation, while ROI 2 exhibited the lowest SSIM (0.446). However, it is important to note that ROI 2 also showed the least pigmentation changes, which may explain its lower SAM and SSIM variability rather than a failure in reconstruction.

**Table 1:** Spectral and Spatial Evaluation using SAM and SSIM across 6 ROIs.

| Metric | ROI 1 | ROI 2 | ROI 3 | ROI 4 | ROI 5 | ROI 6 | Overall |
| --- | --- | --- | --- | --- | --- | --- | --- |
| SAM | $0.453 \pm 0.054$ | $0.610 \pm 0.087$ | $0.466 \pm 0.101$ | $0.531 \pm 0.041$ | $0.488 \pm 0.042$ | $0.568 \pm 0.139$ | $0.519 \pm 0.102$ |
| SSIM | $0.563 \pm 0.024$ | $0.446 \pm 0.036$ | $0.596 \pm 0.019$ | $0.453 \pm 0.031$ | $0.501 \pm 0.008$ | $0.502 \pm 0.008$ | $0.510 \pm 0.059$ |

**Figure 5:**
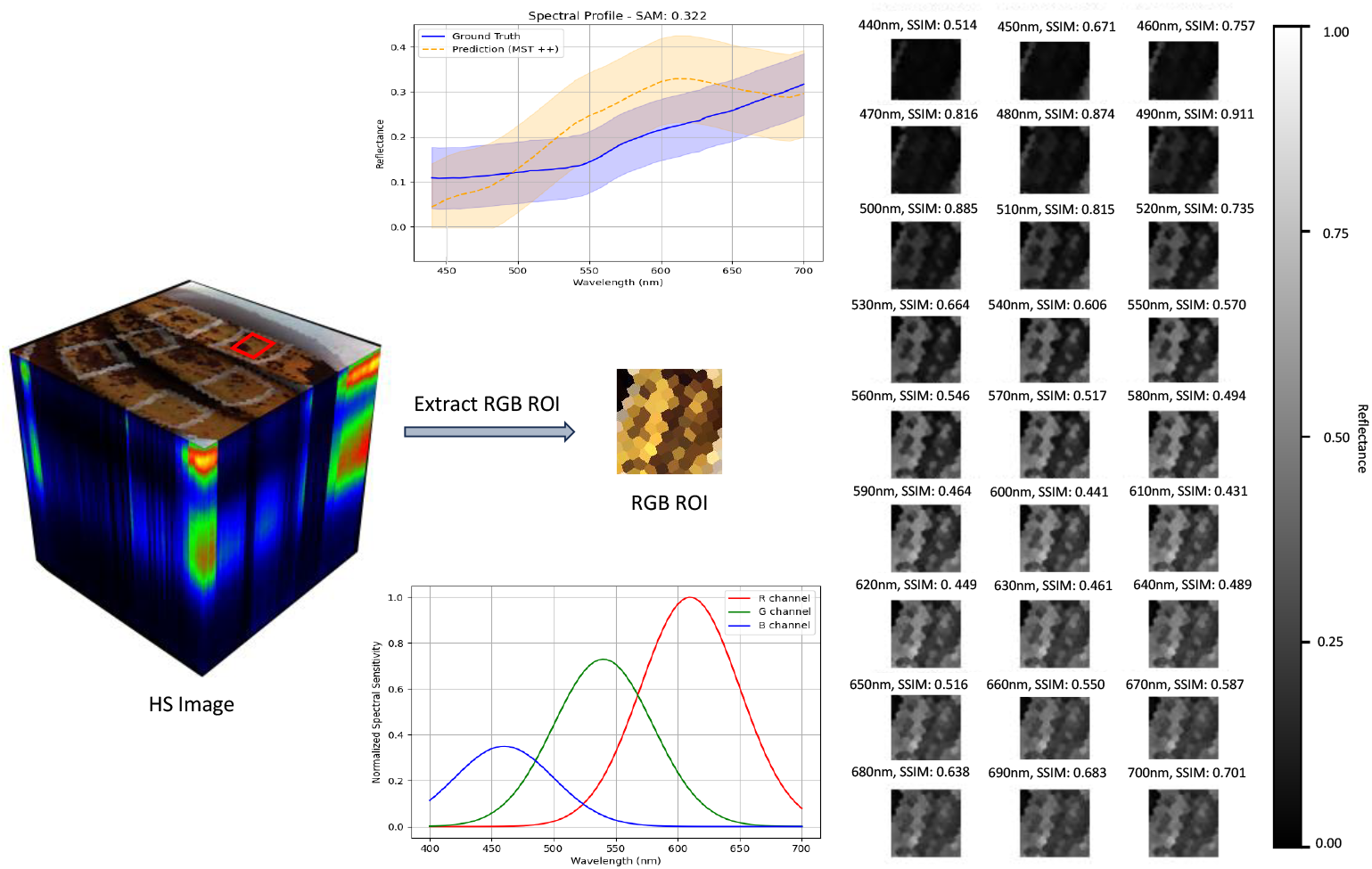
The HS cube and RGB ROI represents the camera response function. A spectral evaluation was performed across 27 image bands to assess the performance of the pre-trained MST++ model. The analysis indicates that the model achieves higher reconstruction accuracy in the mid-range bands (460nm - 520nm) compared to the bands at the two ends. Additionally, spatial assessment reveals that ROIs in these middle bands exhibit improved SSIM relative to the initial bands.

Our dataset demonstrates the feasibility of RGB-to-HSI mapping using the MST++ model, aligning well within its supported spectral range. The evaluation results highlight its effectiveness while revealing spectral variations influenced by oxidative stress and pigment degradation. Although some discrepancies were noted, the findings confirm the potential for accurate spectral reconstruction with further optimisation.

## Benefit and Limitation

This study introduces a non-invasive and cost-effective approach to analyse changes in skin pigmentation using banana peels as a surrogate model, leveraging their biochemical similarity to human skin. The use of HSI enables detailed spectral detecting of pigmentation variations and bruises, providing valuable insights into dermatological conditions such as melasma, vitiligo, and melanoma. Figure 6 demonstrates the advantages of HSI over RGB in tracking pigmentation changes over time for ROI 5. Unlike RGB, which captures only three color channels and misses subtle biochemical variations, HSI provides detailed spectral insights, enabling early detection of pigmentation changes. The ability to track pigmentation changes over time allows early detection of skin abnormalities, which can benefit personalised dermatological treatments and disease monitoring.

**Figure 6:**
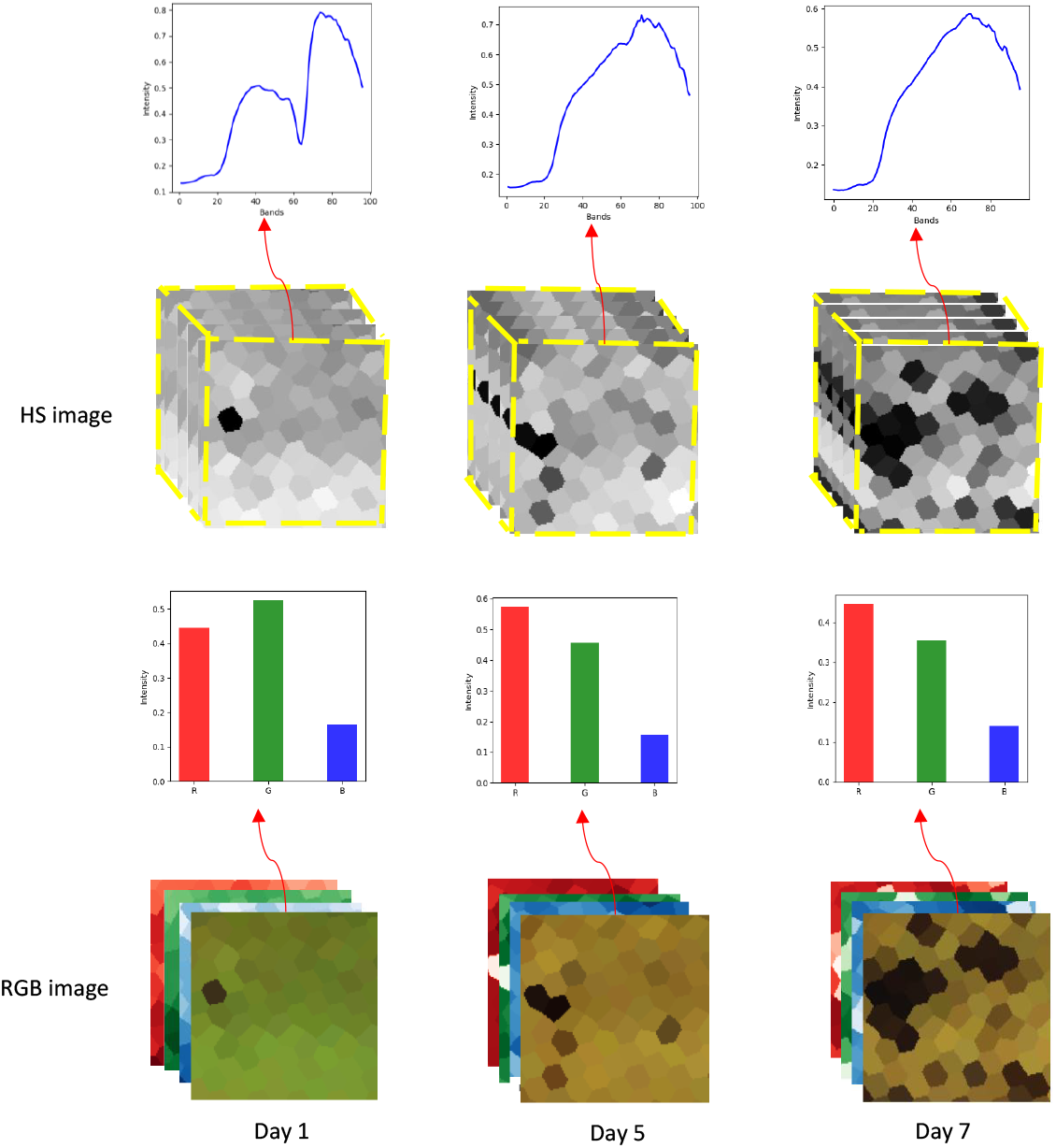
Comparison between RGB and HS Imaging for ROI 5 across Day 1, 4, 7.

Additionally, the study explores RGB-to-HSI mapping, providing a low-cost alternative to direct HSI acquisition, making hyperspectral imaging more accessible for mobile health diagnostics and real-time dermatology applications. The results further validate the effectiveness of hyperspectral reconstruction in detecting pigmentation changes, highlighting the potential for AI-driven skin health monitoring and hyperspectral skin analysis.

However, there are certain limitations. The study was conducted in a natural environment rather than under controlled lighting conditions, leading to potential spectral inconsistencies due to ambient illumination variations. Additionally, RGB-to-HSI mapping requires further optimisation to improve spectral reconstruction accuracy, achieving lower SAM values and higher SSIM scores for enhanced hyperspectral skin imaging. Although banana peels share biochemical similarities with human skin, they do not fully replicate human tissue structure, limiting direct clinical applicability. Future research will focus on controlled lighting conditions, improved spectral reconstruction models, and validation on human skin datasets to enhance dermatological applications. Additionally, further research will also explore human skin characteristics, analysing spectral signatures related to different skin tones, ageing effects, and dermatological conditions to improve practical applications of hyperspectral imaging in dermatology.

## Conclusion

In this study, we introduced an HSI-based imaging approach using banana peels as a surrogate model for detecting pigmentation changes relevant to human skin conditions such as sunspots and melanoma. We developed an HSI protocol that captures spectral reflectance in the 440 nm to 900 nm range, enabling a more precise spectral assessment of biochemical changes over time. Our findings confirm that HSI effectively detects biochemical variations, providing a non-invasive and accurate method for studying pigmentation disorders. To address the computational challenges of direct HSI imaging, we conducted preliminary experiments on RGB-to-HSI mapping, successfully reconstructing 27-band hyperspectral representations in the visible spectrum. The feasibility of AI-driven spectral reconstruction presents a cost-effective alternative for hyperspectral imaging in dermatology.

## Acknowledgements

The work was partially funded by the Centre for Advanced Manufacturing and Materials (CAMM) at the University of Greenwich, within the thematic area in Bio-Inspired Engineering, Imaging & AI (BIO-ENIGMA).

## Author contributions

G.T. and T.N. initialised concepts and directions. N.S. conducted experiments and analysed results. T.N. and G.T. provided critical updates and suggestions that significantly enhanced the scope and direction of the research. N.S. wrote the paper with important input from G.T. and T.N. All authors, N.S., G.T., T.N. reviewed and approved the final manuscript.

## Footnotes

1 https://github.com/livingoptics

